# IgM hyposialylation drives podocyte injury in pediatric and young adult patients with podocytopathies

**DOI:** 10.1101/2025.11.04.686538

**Authors:** Sonia Spinelli, Andrea Garbarino, Francesca Lugani, Edoardo La Porta, Andrea Petretto, Martina Bartolucci, Chiara Lavarello, Nicole Grinovero, Sofia Gaudiano, Ilaria Musante, Paolo Scudieri, Antonella Trivelli, Giorgio Piaggio, Alberto Magnasco, Maria Ludovica Degl’Innocenti, Simona Granata, Gianluigi Zaza, Enrico Verrina, Giovanni Candiano, Maurizio Bruschi

## Abstract

**Introduction:** Altered immunoglobulin glycosylation has been implicated in antibody-mediated podocytopathies, yet the functional impact of IgM sialylation remains unclear. Previous evidence suggested that circulating cationic or hyposialylated IgM may contribute to podocyte injury in idiopathic nephrotic syndrome (iNS).

**Methods:** Serum IgM from pediatric and young adult patients with podocytopathies, membranous nephropathy (MN), lupus nephritis (LN), and healthy controls were analyzed by lectin-based ELISA using a panel of six biotinylated lectins to detect terminal N-glycan residues. Among these, *Sambucus nigra agglutinin* (SNA) binds α2,6-linked sialic acid, and *Ricinus communis agglutinin I* (RCA-I) recognizes β1,4-linked galactose. Serum levels of the α2,6-sialyltransferase ST6GAL1 and the sialidases neuraminidase-1 (NEU1) and neuraminidase-3 (NEU3) were quantified. Cultured human podocytes were exposed to patient-derived, control, or enzymatically modified IgM (desialylated/resialylated) and analyzed by confocal microscopy, quantitative proteomics, phosphoproteomics, and metabolic assays.

**Results:** IgM from patients with podocytopathies showed reduced SNA binding, which inversely correlated with proteinuria (r = −0.69, P < 0.0001) and with serum sialidases NEU1/NEU3 (r = −0.67/−0.59, P < 0.0001). In paired samples, SNA reactivity decreased during relapse and normalized in remission, indicating dynamic modulation with disease activity. ST6GAL1 was undetectable in all groups. In contrast, PLA2R1-positive MN, in which pathogenic antibodies target podocyte PLA2R1, displayed reduced RCA-I binding. Podocytes exposed to hyposialylated or desialylated IgM exhibited actin disorganization, loss of nephrin signal, increased lipid peroxidation, and decreased ATP synthesis. In contrast, resialylated IgM preserved morphology and metabolism comparable to controls. Proteomic and phosphoproteomic profiling revealed enrichment of MAPK, mTOR, AMPK, and cytoskeletal remodeling pathways.

**Conclusions:** IgM hyposialylation, driven by extracellular sialidases up-regulation, correlates with disease activity and promotes oxidative stress, mitochondrial dysfunction, and cytoskeletal remodeling in podocytes, identifying immune glycan remodeling as a modifiable determinant of podocyte injury and a potential biomarker of disease activity.

**Translational Statement:** Altered glycosylation of circulating immunoglobulins is emerging as a key mechanism underlying glomerular injury. Here, we demonstrate that hyposialylated IgM, resulting from elevated serum sialidases NEU1 and NEU3, directly induces podocyte cytoskeletal disorganization, oxidative stress, and metabolic impairment. In contrast, enzymatic resialylation maintains podocyte morphology and function at levels indistinguishable from controls, indicating that sialic acid loss is a modifiable determinant of podocyte injury. Clinically, reduced IgM sialylation correlates with proteinuria and disease activity in idiopathic nephrotic syndrome, supporting its potential as a biomarker of relapse and remission. Targeting the sialidase–sialic acid pathway may therefore represent a promising strategy to preserve podocyte integrity and improve outcomes in antibody-negative podocytopathies.

## INTRODUCTION

Idiopathic nephrotic syndrome (iNS) represents a heterogeneous group of glomerular disorders characterized by podocyte injury, foot-process effacement, and proteinuria. Despite decades of investigation, its immunopathogenesis remains only partially understood. Both clinical and experimental evidence indicate that circulating immune factors can directly alter podocyte structure and function, bridging the historical distinction between immune and non-immune podocytopathies. While the role of IgG autoantibodies has been established in antibody-mediated glomerular diseases such as membranous nephropathy, the contribution of IgM to podocyte injury has received comparatively little attention. Early studies from our group demonstrated that cationic IgM from iNS patients bind to the glomerular basement membrane and can induce proteinuria when injected into rats, supporting a pathogenic role for these antibodies ^1^. More recently, IgM hyposialylation has been identified as a distinctive biochemical signature in pediatric patients with steroid-dependent or frequently relapsing iNS, where hyposialylated IgM persists on T-cell surfaces and alters steroid responsiveness ^2^. These findings suggest that post-translational modifications of immunoglobulins, particularly in their glycan composition, may modulate their immunological behavior and tissue tropism.

Sialic acid is a terminal monosaccharide that confers a negative charge to glycoproteins, modulating molecular interactions, complement activation, and immune recognition. Its loss profoundly affects glomerular permselectivity: mice with mutations in the sialic-acid biosynthetic enzyme GNE (UDP-N-acetylglucosamine 2-epimerase/N-acetylmannosamine kinase) develop severe proteinuria and podocyte effacement, which can be reversed by supplementation with N-acetylmannosamine ^3^. Similarly, reduced sialylation of podocyte podocalyxin or secreted angiopoietin-like 4 (ANGPTL4) alters glomerular charge and induces proteinuria ^4^. In this context, sialylation functions not merely as a structural modification but as a regulatory checkpoint of both immune and non-immune mediated podocyte integrity.

Beyond sialylation, other glycan alterations, such as IgG afucosylation, have recently been linked to antibody-mediated kidney injury. In patients with iNS harboring anti-nephrin autoantibodies, circulating and urinary IgG displayed markedly reduced fucosylation, correlating with disease activity and proteinuria ^5^. Together, these data highlight that immune glycan remodeling, through altered fucosylation or sialylation, may fundamentally shape antibody effector functions and podocyte interactions.

In the present study, we investigated whether hyposialylation contributes directly to podocyte injury and metabolic dysfunction. We combined lectin-based glycoprofiling of IgM from patients with podocytopathies with functional assays in cultured human podocytes exposed to native, desialylated, or resialylated IgM. We also quantified sialidase (NEU1/NEU3) levels in patient sera to identify potential enzymatic drivers of IgM desialylation. Our results support that IgM hyposialylation promotes MAPK activation, oxidative stress, and ATP depletion in podocytes, and that these alterations are reversible upon resialylation, establishing reduced sialylation as a modifiable determinant of podocyte injury.

## METHODS

Serum IgM from patients with podocytopathies, membranous nephropathy, lupus nephritis, and healthy controls was analyzed by lectin-based ELISA to assess N-glycan residues. Serum levels of sialylation- and desialylation-related enzymes (ST6GAL1, NEU1, and NEU3) were quantified by direct ELISA. Purified IgM from selected patients and controls, as well as enzymatically desialylated and resialylated IgM, were used to treat cultured human podocytes for confocal imaging, quantitative proteomic and phosphoproteomic profiling, and metabolic assays of oxidative stress and ATP synthesis. Detailed experimental procedures, including patient selection criteria ^5, 6^, IgM purification and enzymatic modification protocols, lectin- and enzyme-ELISA conditions ^7, 8^, cell culture ^9, 10^, confocal microscopy ^11^, proteomic workflow ^12, 13^, kinase array analysis ^14^, and statistical methods ^14-16^, are provided in the Supplementary Methods.

## RESULTS

### Clinical and biochemical characteristics of the study population

Serum samples were obtained from 56 pediatric and young adult patients with biopsy-proven glomerular disease, including 32 with minimal change disease (MCD) and 24 with focal segmental glomerulosclerosis (FSGS). As shown in Table 1, patients displayed variable degrees of proteinuria, ranging from normal to overt nephrotic levels. They were characterized by different clinical phenotypes (steroid-dependent, multidrug-dependent, or multidrug-resistant). None of the MCD or FSGS patients were positive for circulating anti-nephrin (NPHS1).

**Table 1.** Clinical and demographic characteristics of the study cohort. Summary of demographic, clinical, and histopathological features of patients with podocytopathies [minimal change disease (MCD), focal segmental glomerulosclerosis (FSGS)], membranous nephropathy (MN) (including PLA2R1-positive and -negative cases), lupus nephritis (LN), and healthy controls (CTR). Values are expressed as median (interquartile range) or number (percentage) as appropriate.

|  | FSGS | MCD | MN | LN |
| --- | --- | --- | --- | --- |
| Number (%) | 24(43) | 32(57) | 20 | 20 |
| Female (%) | 10(42) | 14(44) | 10(50) | 10(50) |
| Male (%) | 14(58) | 18(56) | 10(50) | 10(50) |
| Age median (range) years | 7(2-21) | 8(3-21) | 45(31-51) | 39(28-49) |
| UPCR mg/mg | 4(0.05-13) | 3.5(0.05-9) | 4(1.1-13) | 3(0.5-7) |
| Serum anti-Nephrin positive | 0 | 0 | 0 | 0 |
| Urine anti-Nephrin positive | 0 | 0 | 0 | 0 |
| Serum anti-PLA2R positive | Not tested | Not tested | 10 | Not tested |

### Lectin-based profiling of N-glycan residues on serum IgM

Lectin specificity was verified by pretreating purified serum IgM (Figure S1A) with N-glycosidase F (PNGase F), which completely abolished lectin binding (Figure S1B), confirming that the signals reflected N-glycan–dependent recognition.

The glycosylation profile of serum IgM was assessed by lectin ELISA using six biotinylated lectins: Sambucus nigra agglutinin (SNA), Aleuria aurantia lectin (AAL), Ulex europaeus agglutinin I (UEA-I), Lotus tetragonolobus lectin (LTL), Ricinus communis agglutinin I (RCA-I), and Concanavalin A (ConA). Each lectin preferentially binds specific glycan motifs: SNA to terminal α2,6-linked sialic acid (Neu5Acα2-6Gal/GalNAc), AAL to fucose, UEA-I to α1,2-linked fucose, LTL to α-linked fucose or mannose, RCA-I to β1,4-linked galactose, and ConA to mannose or hybrid-type residues.

As shown in Figure 1A, binding to SNA was markedly reduced in patients with podocytopathies (499[480-527] RU/ml) compared with healthy controls (CTR; 614[554-654] RU/ml), lupus nephritis (LN; 583[555-612] RU/ml), and membranous nephropathy (MN; 581[568-591] RU/ml) (P < 0.0001).

**Figure 1.**
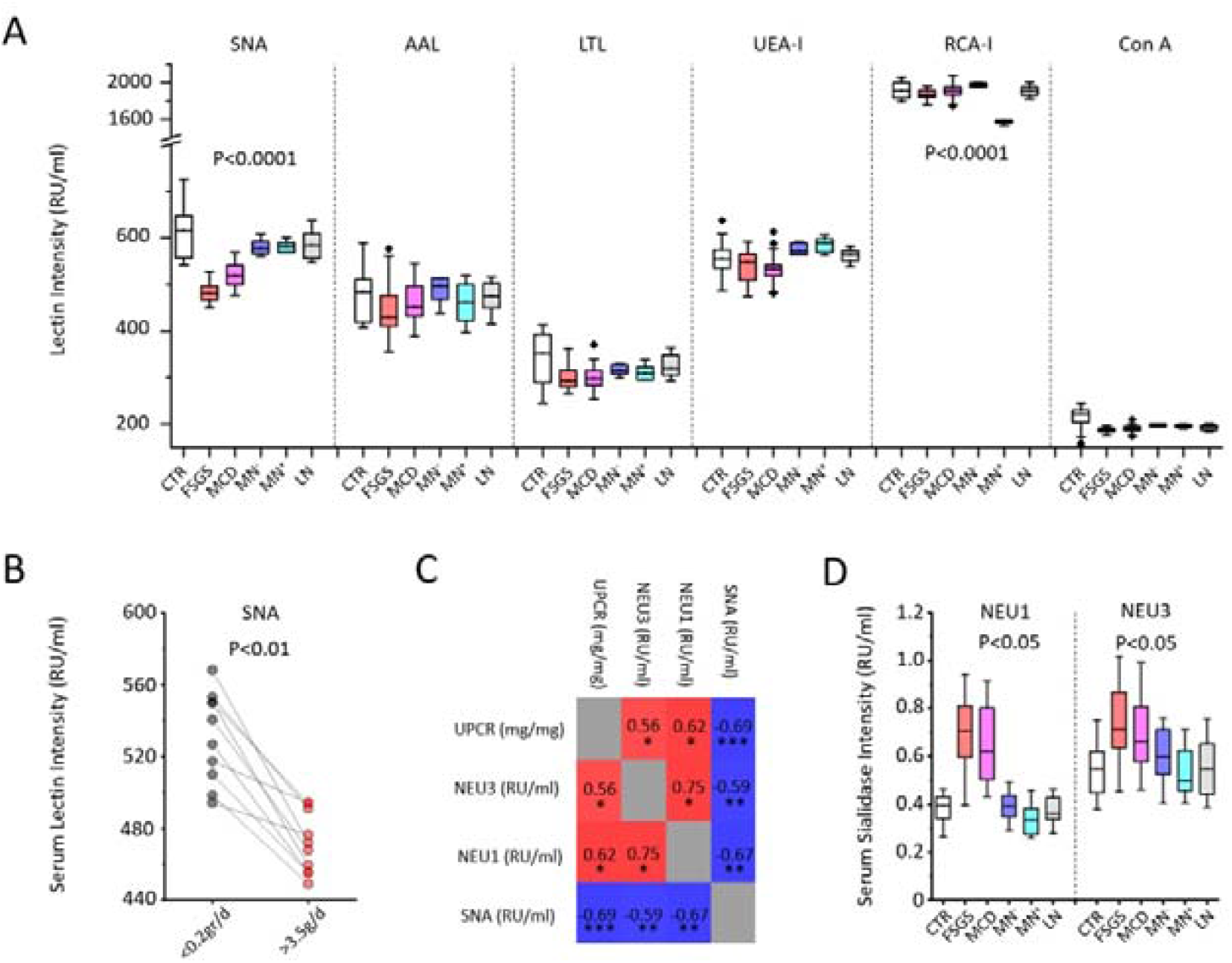
Lectin-based IgM N-glycan profiling and serum neuraminidase levels. (A) Boxplots showing lectin-ELISA results for IgM N-glycan residues across control (CTR; white), minimal change disease (MCD; magenta), focal segmental glomerulosclerosis (FSGS; red), membranous nephropathy PLA2R1 negative (MN^-^; blue) or positive (MN^+^; cyan), and lupus nephritis (LN; gray). Binding to SNA was significantly reduced in FSGS and MCD (P < 0.0001), while RCA-I binding was lowest in PLA2R1-positive MN (P < 0.0001). (B) Paired analysis of SNA reactivity in ten patients with podocytopathies at two clinical time points, during active disease (proteinuria > 3.5 g/day) and complete remission (proteinuria < 0.2 g/day). SNA binding was significantly reduced during the active phase compared with remission (P < 0.01). (C) Heatmap correlogram displaying Pearson correlation coefficients among SNA reactivity, NEU1 and NEU3 serum levels, and urinary protein-to-creatinine ratio (UPCR). The color scale indicates the strength and direction of the correlation: red represents a strong positive, blue a strong negative, and white no correlation; color intensity is proportional to the absolute value of the correlation coefficient (*** = P<0001; ** = P<0.001; * = P<0.05). (D) Boxplots of NEU1 and NEU3 serum concentrations showing significant increases in podocytopathies versus other groups (P < 0.001). ST6GAL1 was undetectable in all samples (data not shown).

Among podocytopathies, patients with FSGS (479[466-496] RU/ml) showed significantly lower SNA reactivity than those with MCD (517[499-540] RU/ml) (P < 0.0001), indicating a more pronounced loss of terminal sialic acid in the FSGS subgroup.

All lectin signals were normalized to the total IgM content of each sample, ensuring that the observed differences reflected glycosylation rather than concentration variability.

Notably, in ten patients with podocytopathies for whom paired serum samples were available at two distinct clinical time points, during active disease (proteinuria > 3.5 g/day; SNA = 470[455-492]) and complete remission (proteinuria < 0.2 g/day; SNA = 534[5075-551]), SNA binding was significantly reduced during the active phase compared with remission (P < 0.01), confirming that IgM sialylation dynamically reflects disease activity (Figure 1B).

Furthermore, Pearson correlation analysis showed that IgM hyposialylation correlated inversely with circulating NEU1 (R = –0.67, P < 0.001) and NEU3 (R = –0.59, P < 0.001) levels, and with the urinary protein-to-creatinine ratio (UPCR) (R = –0.69, P < 0.0001) (Figure 1C).

For all other lectins tested (AAL, LTL, UEA-I, and ConA), no significant differences were observed among groups.

In contrast, RCA-I, which recognizes terminal galactose residues with a preference for β1,4 linkages, revealed a distinct pattern (Figure 1A): MN patients positive for circulating anti-PLA2R1 antibodies (1581[1555-1594]) displayed markedly reduced RCA-I binding compared with all other groups, including PLA2R1-negative MN (1978[1953-1998]), LN (1915[1863-1953]), iNS (1885[1856-1937]), and CTR (1911[1827-2002]) (P < 0.0001).

### Serum Expression levels of sialylation- and desialylation-related enzymes in serum

To investigate whether the reduced sialic acid levels observed on IgM could be attributed to altered sialylation or increased desialylation, a direct ELISA was performed for ST6 beta-galactoside alpha-2,6-sialyltransferase 1 (ST6GAL1), neuraminidase 1 (NEU1), and neuraminidase 3 (NEU3) in serum samples from all study groups. ST6GAL1 levels were undetectable in all samples analyzed (data not shown). In contrast, both NEU1 and NEU3 were significantly higher in patients with iNS (NEU1 = 0.65[0.53-0.81]; NEU3 = 0.69[0.59-0.85]) compared with CTR (NEU1 = 0.39[0.34-0.43]; NEU3 = 0.548[0.45-0.628]), LN (NEU1 = 0.36[0.33-0.43]; NEU3 = 0.549[0.44-0.66]), and MN (NEU1 = 0.366[0.32-0.41]; NEU3 = 0.569[0.46-0.69]) (P<0.0001 for both enzymes). Notably, among podocytopathies, FSGS patients exhibited significantly higher serum NEU1 and NEU3 levels (NEU1 = 0.71[0.59-0.81]; NEU3 = 0.72[0.63-0.88]) than those with MCD (NEU1 = 0.62[0.50-0.80]; NEU3 = 0.67[0.58-0.81]) (P < 0.05) (Figure 1D), supporting a link between increased sialidase abundance and the severity of IgM hyposialylation of different study groups.

No significant differences in NEU1 or NEU3 were observed between LN and MN compared to CTR. A significant positive Pearson correlation was found between NEU1 and NEU3 concentrations across all samples (R = 0.75, P < 0.05), with NEU3 levels consistently higher than NEU1 in all study groups.

### Effects of IgM sialylation status on podocyte cytoskeletal organization and nephrin expression profile

As shown in Figure S1B, the efficiency of enzymatic desialylation and subsequent resialylation was validated by SNA lectin ELISA, whereas, to exclude residual sialyltransferase or sialidase activity in the IgM preparations used for podocyte stimulation, desialylated and resialylated IgM were tested by specific ELISA against ST6GAL1, NEU1, and NEU3 (Figure S1C). All results were negative, confirming the absence of enzyme carryover.

To determine whether IgM purified from patients with podocytopathies could directly induce structural alterations, immortalized human podocytes were incubated with IgM purified from five independent iNS pools or from matched CTR pools, both processed under identical conditions. Each iNS pool included sera from four patients with low but comparable sialic acid levels, ensuring consistency across experiments.

Preliminary experiments testing different IgM concentrations and incubation times revealed a direct correlation between both parameters and the extent of cytoskeletal rearrangement (Figure S2). Based on these optimization results, a concentration of 0.12 µg IgM per well in PBS (corresponding to a 1:50 serum dilution) incubated overnight was selected as the standard experimental condition, as it produced the maximal and most reproducible effect. As shown in Figures 2A and 2C, untreated podocytes and those exposed to CTR IgM exhibited comparable morphology and actin organization. In contrast, iNS IgM caused prominent cytoskeletal rearrangement, with reduced phalloidin fluorescence, cell shrinkage, and a marked decrease in adhesion (P < 0.0001). Desialylated CTR IgM reproduced and intensified these effects, inducing severe actin disorganization and stress-fiber loss. By comparison, resialylated IgM did not alter podocyte morphology or adhesion, yielding actin patterns and cell counts comparable to those observed with control IgM.

**Figure 2.**
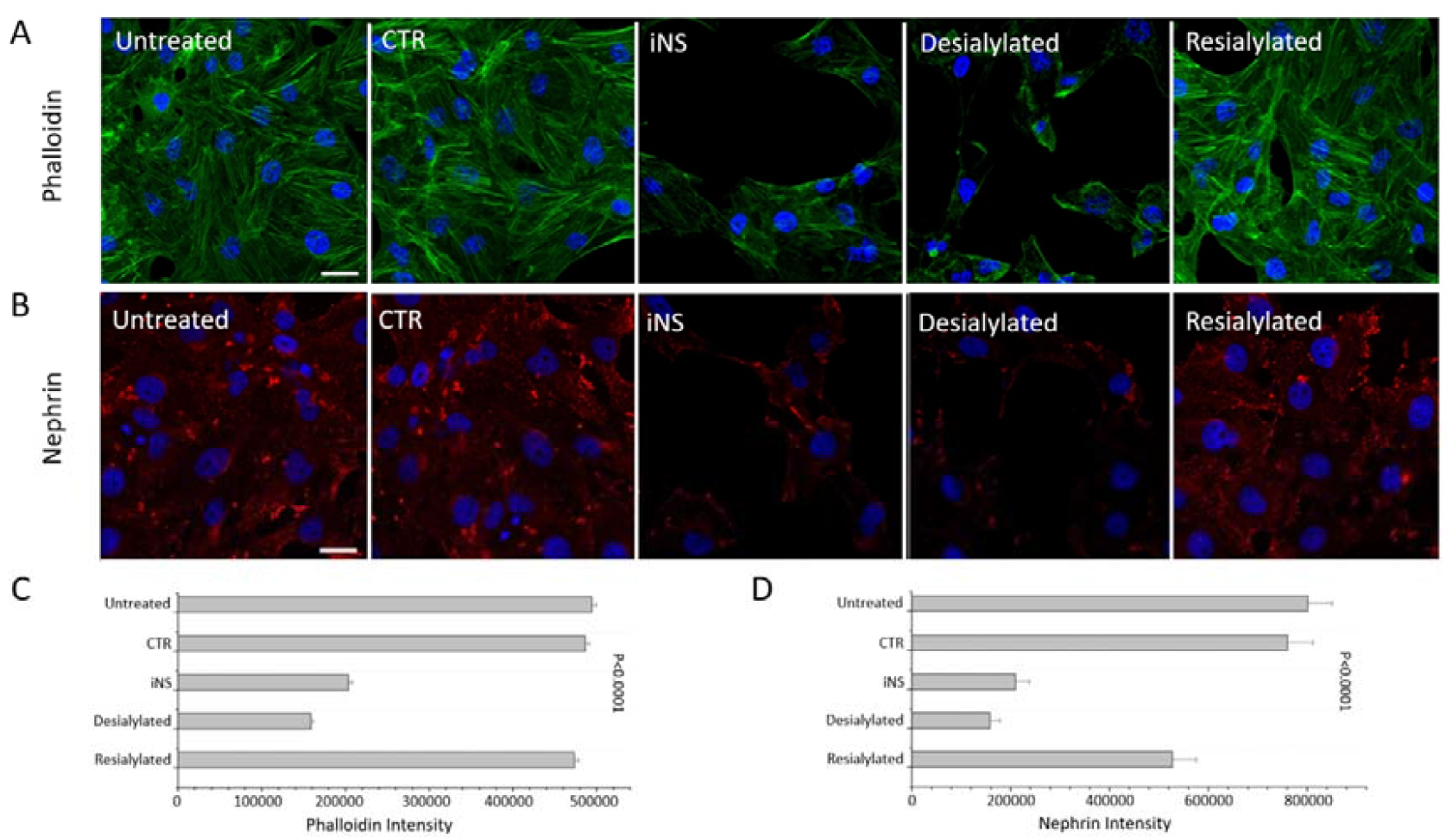
Confocal immunofluorescence analysis of podocytes exposed to IgM with different sialylation status. (A–B) Representative images showing phalloidin-stained actin cytoskeleton (green) and nephrin distribution (red); nuclei in blue. Untreated and CTR IgM–treated podocytes display normal morphology with preserved stress fibers and intense nephrin staining. In contrast, iNS-derived and desialylated IgM induce actin disorganization, loss of stress fibers, cell shrinkage, reduced adhesion, and a marked decrease in nephrin signal intensity. Resialylated IgM maintains actin architecture and nephrin localization to levels indistinguishable from control cells. (Scale bars: 10 µm.). (C–D) Quantitative analysis corresponding to panels (A–B), showing total phalloidin (C) and nephrin (D) fluorescence per field, computed with EBImage (sum of pixel intensities under identical acquisition settings). The quantitative results confirm the reduction in actin and nephrin signals in iNS- and desialylated IgM–treated podocytes, and their recovery after resialylation.

Consistent results were obtained for nephrin immunostaining: CTR and resialylated IgM preserved normal nephrin distribution, whereas iNS and desialylated IgM reduced signal intensity and disrupted membrane localization (P < 0.0001; Figure 2B, D).

### Podocyte proteome profile

Label-free quantitative proteomic analysis identified 6,159 proteins across all samples. Principal component analysis of the whole proteome dataset revealed a clear separation among the three experimental conditions, podocytes treated with CTR IgM, iNS IgM, or desialylated IgM, indicating distinct global protein expression patterns (Figure 3A).

**Figure 3.**
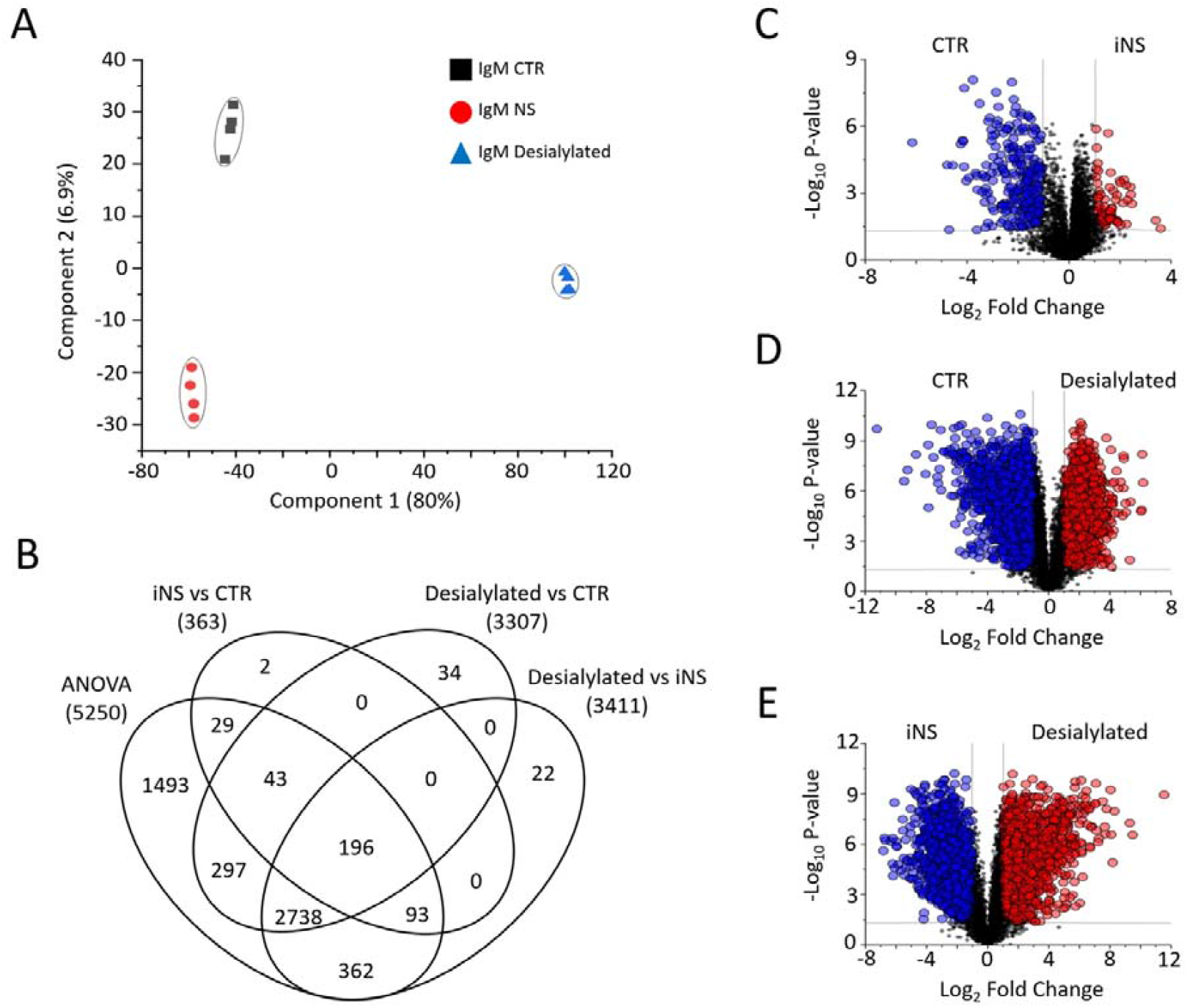
Proteomic analysis of podocytes exposed to IgM of different sialylation states. (A) Principal component analysis (PCA) showing distinct clustering of podocytes treated with CTR, iNS, or desialylated IgM. (B) Venn diagram summarizing the overlap of statistically significant proteins identified by ANOVA and unpaired t-tests. Numbers and circles represent the distinct statistically significant proteins in each comparison, respectively. (C–E) Volcano plots comparing (C) iNS vs CTR, (D) Desialylated vs CTR, and (E) Desialylated vs iNS. In the volcano plot, the x-axis reports log_2_ fold change, and the y-axis: –log_10_ P-value. Black dots indicate non-significant proteins; red and blue denote significantly up- or down-regulated proteins, respectively.

An ANOVA test for unpaired samples identified 5,250 proteins showing significant variation among groups (Table S1, Figure 3B). To further dissect these differences, unpaired t-tests were performed. The comparison between iNS and CTR IgM-treated cells revealed moderate changes in 363 proteins, suggesting a defined yet relatively contained alteration in podocyte homeostasis. In contrast, exposure to desialylated IgM led to a much broader proteomic response, with 3,037 proteins differing from CTR IgM-treated cells. When desialylated IgM was directly compared with iNS IgM, 3,411 proteins were found to be significantly modulated, underscoring the substantial impact of complete sialic acid removal on podocyte protein networks (Table S1, Figure 3B-E).

Altogether, these data are consistent with a significant role for IgM sialylation in proteome remodeling.

### Gene Ontology enrichment analysis

To elucidate the biological significance of the proteomic changes observed, we performed Gene Ontology (GO) enrichment analysis of the sets of statistically significant proteins identified across the three comparisons. This analysis revealed 38 significantly enriched GO terms (Table S2) that could be grouped into five major biological clusters: MAPK signaling and stress response, cell–cell and cell–matrix interactions, mitochondrial dysfunction and metabolic adaptation, oxidative stress, and inflammatory signaling (Figure 4).

**Figure 4.**
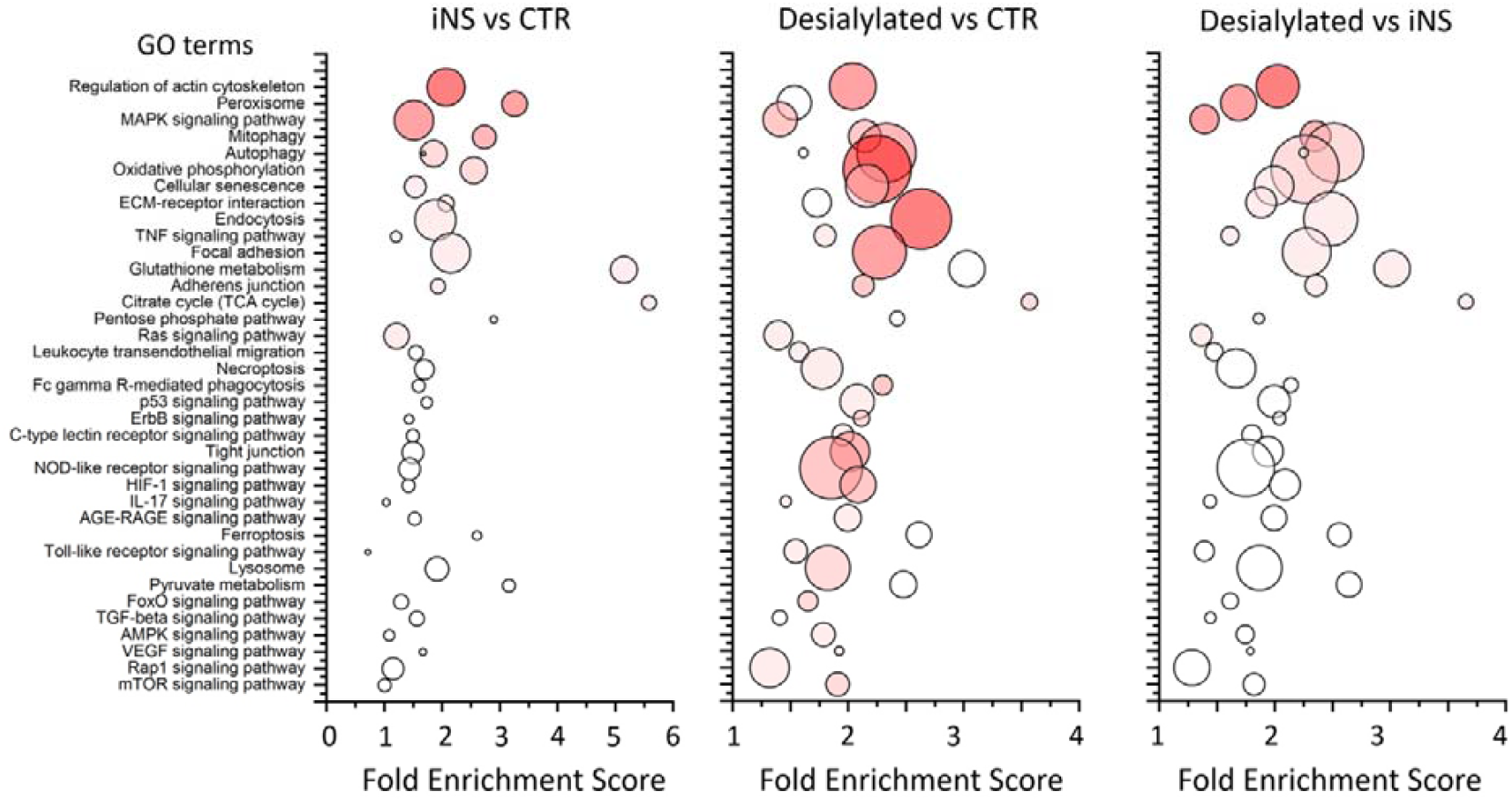
Gene Ontology (GO) enrichment analysis of differentially expressed proteins. Each circle represents a significantly enriched GO Biological Process term. The x-axis indicates the Fold Enrichment Score; circle size is proportional to the number of associated proteins, and color intensity reflects statistical significance (white = –log_10_ 1.3; red = – log_10_ 38). Major clusters include MAPK and stress-response signaling, cytoskeletal organization, mitochondrial and metabolic pathways, oxidative stress, and inflammatory signaling.

Specifically, pathways related to MAPK, Ras, mTOR, and AMPK signaling were among the most significantly modulated, indicating a broad remodeling of intracellular signaling cascades. Processes involved in cytoskeletal dynamics and adhesion, such as the regulation of the actin cytoskeleton, focal adhesion, ECM–receptor interaction, adherence junctions, and tight junctions, were also prominently affected, in line with the morphological alterations observed by microscopy. The concomitant enrichment of oxidative phosphorylation, TCA cycle, peroxisome, and glutathione metabolism pathways indicated a state of mitochondrial stress and redox imbalance. At the same time, the activation of autophagy, mitophagy, cellular senescence, and ferroptosis suggested the engagement of adaptive or degenerative stress responses.

Finally, multiple immune- and inflammation-related pathways, including TNF, IL-17, Toll-like receptors, NOD-like receptors, HIF-1, TGF-β, and AGE–RAGE signaling, were enriched across all conditions, supporting the concept that loss of IgM sialylation promotes a proinflammatory and stress-related phenotype in podocytes.

Remarkably, these 38 GO terms were consistently detected across all three comparisons, differing only in the degree of enrichment, P-value significance, and the number of associated proteins, suggesting a shared core of perturbed biological processes modulated in intensity by the sialylation status of IgM.

### Phosphoproteomic and kinase activity analysis

To validate and extend the proteomic findings (Figure 5A), a phosphoproteomic profiling was performed on podocytes exposed to IgM purified from CTR or iNS pools, as well as from enzymatically desialylated or reasilylated CTR IgM. Quantitative analysis revealed distinct phosphorylation signatures among conditions, consistent with the proteomic results (Figure 5B).

**Figure 5.**
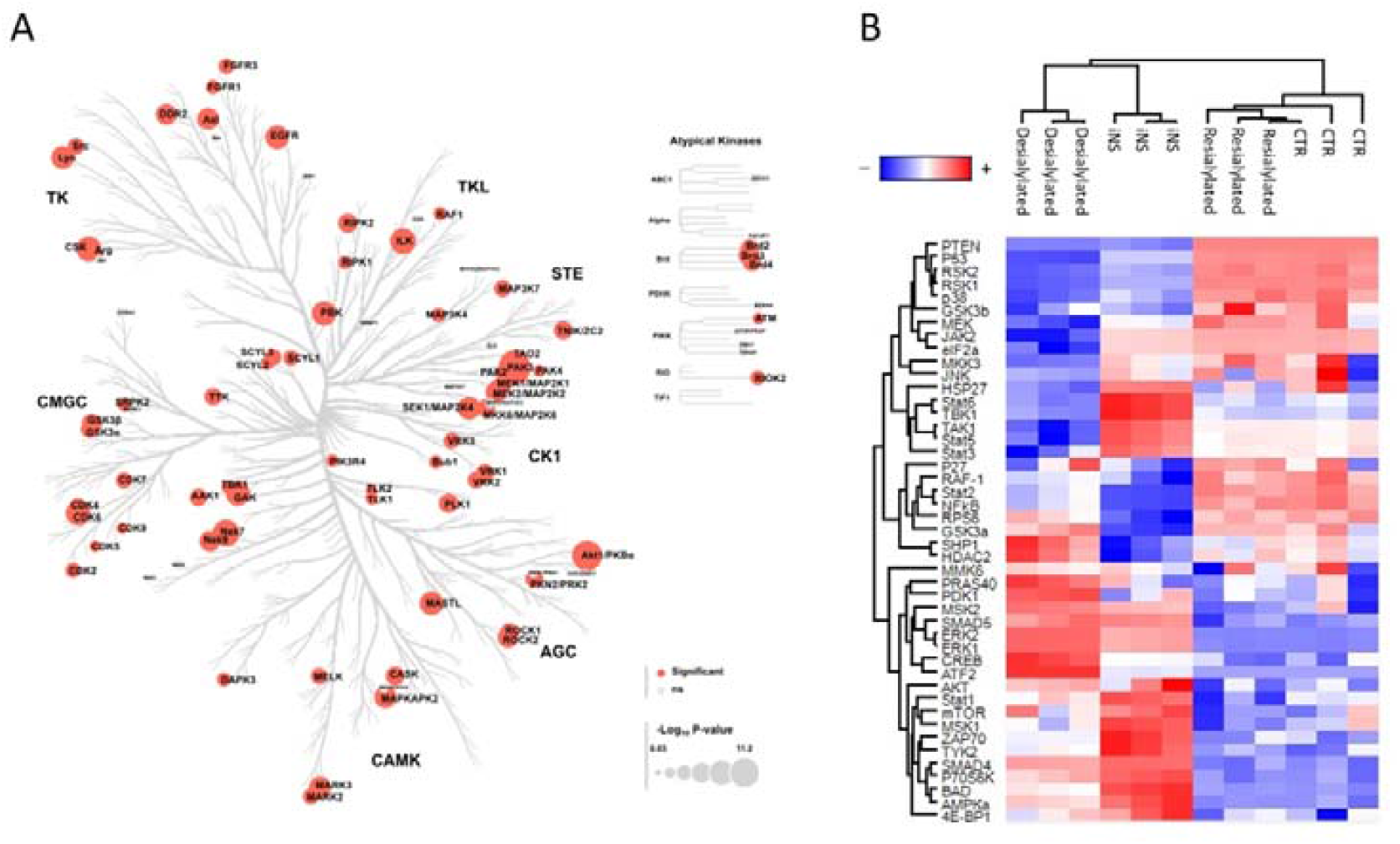
Phospho-array and kinase activity profiling of podocytes exposed to IgM with different sialylation status. (A) The entire kinase family identified by mass spectrometry in podocytes treated with IgM derived from iNS, control (CTR), desialylated, or resialylated preparations is shown. Each colored node represents an individual kinase whose change in expression among the four conditions is either statistically significant (red) or not (gray). Node size is proportional to the –log10 P-value. (B) Heatmap of phospho-array signals showing relative phosphorylation levels in podocytes treated with control (CTR), iNS, Desialylated, and Resialylated CTR IgM. Color intensity represents normalized signal intensity for each phospho-site. In the heatmap, the rows and columns correspond to the kinases and the experimental conditions, respectively.

Kinase-Substrate Enrichment Analysis (KSEA) indicated selective activation of MAPK-related kinases (ERK1/2, p38, JNK) and reduced AKT1–mTOR signaling in iNS-treated cells, reflecting a stress-responsive phenotype. Desialylated IgM further amplified this pattern, with strong activation of MAPK14, MAPK8, PRKCA, and SRC, and inhibition of AKT1, GSK3β, and PAK1, consistent with cytoskeletal remodeling and proinflammatory signaling.

Notably, resialylated IgM did not alter the kinase activity profile, which remained indistinguishable from controls. Together, these findings indicate that loss of IgM sialic acid enhances stress signaling and suppresses survival pathways, whereas resialylation preserves the physiological kinase balance (Figure S4).

### Hyposialylated IgM increases lipid peroxidation and reduces ATP synthesis in podocytes

To assess whether the signaling alterations induced by hyposialylated IgM translated into functional metabolic consequences, we measured lipid peroxidation and ATP synthesis in cultured podocytes exposed to IgM with different sialylation status. Cells were treated with IgM purified from healthy donors (CTR), from patients with podocytopathies (iNS), or with the same CTR IgM preparations after enzymatic desialylation or resialylation. All IgM preparations were processed under identical conditions, extensively washed using Amicon Ultra filters (100 kDa cut-off), and filtered through 0.45 μm membranes prior to cell exposure to ensure the absence of residual enzymes or contaminants. As shown in Figure 6A, levels of malondialdehyde (MDA), a marker of lipid peroxidation, were markedly increased in podocytes treated with desialylated IgM compared with all other groups (P < 0.0001). In contrast, MDA concentrations in cells exposed to resialylated IgM remained at levels indistinguishable from those observed with CTR IgM, confirming that the presence of sialic acid residues prevents oxidative membrane damage. A complementary trend was observed for ATP synthesis: resialylated IgM maintained ATP levels comparable to CTR (Figure 6B). Podocytes treated with desialylated IgM displayed a pronounced reduction in ATP production, indicating mitochondrial dysfunction. Intermediate ATP levels were detected following exposure to iNS IgM, while resialylated IgM preserved ATP generation at control-like values (P < 0.0001). These results were consistent across biological replicates derived from five independent iNS IgM pools, confirming that the observed metabolic effects were pool-independent and strictly dependent on the degree of IgM sialylation.

**Figure 6.**
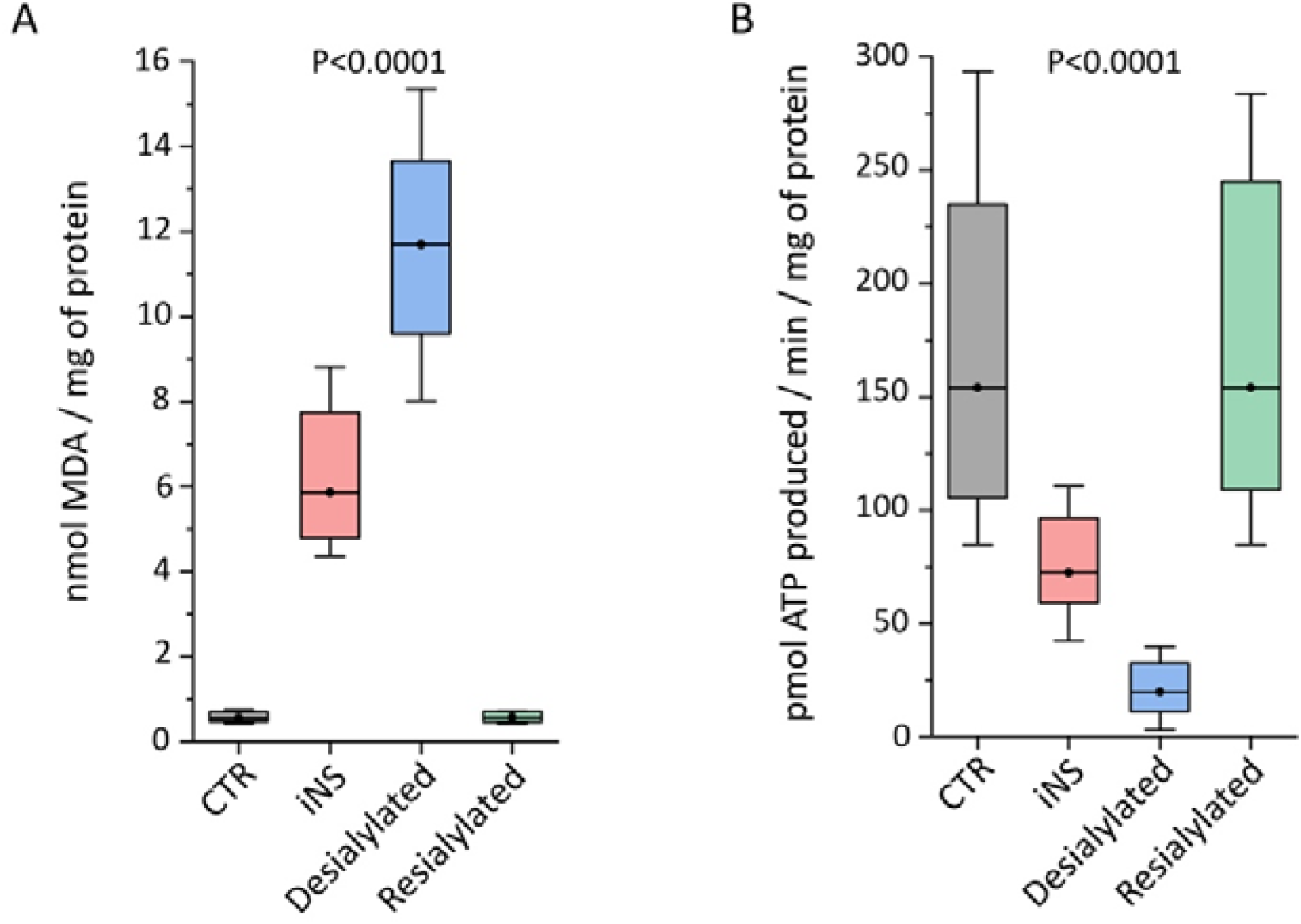
Effects of IgM sialylation status on oxidative stress and ATP production. (A) Boxplots of malondialdehyde (MDA) concentrations showing increased lipid peroxidation in podocytes exposed to iNS-derived IgM and desialylated CTR IgM compared with CTR-derived and resialylated IgM (P < 0.0001). (B) Boxplots of ATP synthesis showing a parallel reduction in cellular ATP levels in iNS-derived and desialylated IgM–treated cells, whereas resialylated IgM maintained ATP levels comparable to control conditions (P < 0.0001). Values are expressed as nmol/mg total protein; each condition was analyzed in ten independent replicates.

Together, these findings are consistent with IgM desialylation driving oxidative stress and decreasing bulk ATP levels in podocytes in vitro. Both parameters returned to baseline levels following IgM resialylation, confirming the functional reversibility of the metabolic alterations associated with sialic acid loss. These metabolic findings, in line with the proteomic and phosphoproteomic data, further support that IgM hyposialylation amplifies stress-driven signaling pathways that compromise podocyte homeostasis.

## DISCUSSION

In this study, we identify IgM hyposialylation as a modifiable biochemical determinant of podocyte injury. Patient-derived IgM with reduced SNA reactivity, and more markedly, enzymatically desialylated IgM, induced actin disorganization, loss of stress fibers, reduced adhesion, decreased nephrin expression, increased lipid peroxidation, and diminished ATP synthesis. Conversely, resialylation maintained structural and metabolic features at levels comparable to healthy controls.

Together with the elevated serum NEU1 and NEU3 detected in the same patients, these data point to a mechanism in which enhanced desialylation of circulating IgM promotes a stress-responsive, energetically compromised podocyte state. Our findings extend and mechanistically connect two prior strands of evidence. First, IgM as pathogenic effectors in podocytopathies: Musante et al. demonstrated that anti-actin/β-ATP-synthase IgM from idiopathic nephrotic syndrome (iNS) patients was cationic, deposited in glomeruli, and induced proteinuria in rats following injection, defining a direct antibody-mediated mechanism of podocyte injury ^1^. Second, IgM hyposialylation on T cells identifies a severe steroid-dependent iNS phenotype: Colucci et al. showed that less-sialylated IgM accumulates on T-cell surfaces, resists internalization, abrogates steroid-induced inhibition, and triggers the release of podocyte-damaging factors, with total IgM sialylation inversely correlating with T-cell–bound IgM ^2^. Our data unify these observations by showing that the sialylation state per se modulates the podocyte-directed toxicity of IgM in vitro.

The central role of sialic acid metabolism in glomerular perm-selectivity is well established. Genetic deficiency of UDP-N-acetylglucosamine 2-epimerase/N-acetylmannosamine kinase (GNE), the key enzyme in sialic-acid biosynthesis, causes severe proteinuria in mice and is rescued by N-acetylmannosamine supplementation, proving that deficient sialylation is sufficient to disrupt the filtration barrier ^3^. Selective desialylation of podocyte glycoproteins, such as podocalyxin, also causes foot-process effacement and FSGS-like lesions ^17, 18^. Furthermore, changes in the sialylation of angiopoietin-like-4 (ANGPTL4) modulate its isoelectric point and glomerular charge selectivity, correlating with proteinuria in steroid-sensitive models ^4^.

At the signaling level, our phosphoproteomic and KSEA analyses revealed a consistent activation of the MAPK (ERK, p38, JNK) cascade and a concomitant down-regulation of AKT/mTOR, consistent with a stress-activated, pro-apoptotic signature expected from NEU-driven desialylation. Indeed, NEU3, a plasma-membrane sialidase, removes sialic acids from gangliosides and modulates β1-integrin trafficking and EGFR activation, thereby altering adhesion and MAPK/PI3K signaling ^19^. Similarly, NEU1, the major lysosomal neuraminidase, regulates growth-factor receptor desialylation and renal homeostasis ^20^. The concurrent elevation of NEU1 and NEU3 in patient sera therefore provides a plausible enzymatic basis for IgM hyposialylation and podocyte dysfunction.

Recent studies have demonstrated that NEU1 and NEU3 are not merely passive lysosomal or membrane-associated enzymes but are dynamically upregulated during inflammation, contributing directly to the desialylation of extracellular and membrane glycoproteins. In idiopathic pulmonary fibrosis (IPF), NEU3 expression and activity are markedly increased in fibrotic lesions and bronchoalveolar lavage fluid, paralleling a global reduction of α2,6-linked sialic acid residues and enhanced SNA reactivity on tissue proteins; pharmacologic or genetic inhibition of NEU3 attenuates inflammation and collagen deposition ^21^. Recombinant NEU3 instillation is sufficient to induce pulmonary inflammation and fibrosis in vivo, confirming its causal role in inflammatory desialylation ^22^. Likewise, NEU1 expression and enzymatic activity are upregulated in activated microglia and lipopolysaccharide-induced neuroinflammation, leading to desialylation of neuronal surface glycoproteins and amplification of oxidative and metabolic stress ^23^. In airway epithelial cells, NEU1 directly removes terminal sialic acids from MUC1 and the EGF receptor, enhancing pro-inflammatory signaling ^24^. Consistently, increased NEU3 expression was also implicated in experimental colitis, where its activity drives mucosal injury and immune activation ^25^. Collectively, these findings support an inflammation-driven axis of neuraminidase upregulation and protein desialylation that parallels the pattern observed in our patients, providing a mechanistic bridge between systemic inflammatory activation, altered serum neuraminidase levels, and IgM hyposialylation–dependent podocyte injury.

Although enzyme activity was not directly measured, the inverse correlation between serum NEU1/NEU3 abundance and IgM sialylation, together with the direct association with proteinuria, strongly suggests that increased neuraminidase activity contributes to antibody desialylation in vivo. This interpretation is further supported by longitudinal analysis of paired samples from patients with MCD or FSGS, showing a significant reduction in SNA reactivity during active disease and a return to normal levels upon remission, demonstrating that IgM sialylation is a reversible, activity-dependent feature. Although ST6GAL1, the main sialyltransferase, was not detected in serum, this finding is consistent with its predominant intracellular localization in the Golgi apparatus rather than secretion into the circulation. It therefore does not contradict a systemic imbalance favoring desialylation. Importantly, this association is consistent with recent evidence that inflammatory stimuli drive transcriptional and functional upregulation of NEU1 and NEU3, leading to widespread desialylation of membrane and secreted glycoproteins in fibrotic and autoimmune settings ^21, 23^. The inflammatory environment typical of iNS may favor systemic sialidase activation, resulting in IgM hyposialylation and increased podocyte vulnerability. In functional assays, desialylated IgM recapitulated and intensified the podocyte alterations observed with patient IgM, whereas resialylated IgM maintained physiological cytoskeletal and metabolic integrity comparable to that of controls. This model supports a causal role for sialylation status in podocyte responses in vitro and fits within a broader paradigm in which inflammation-induced sialidase activation alters protein sialylation and cellular homeostasis. It also aligns with classical evidence that cationic or desialylated proteins cross the glomerular basement membrane more readily and induce proteinuria ^26^. These findings provide a mechanistic framework connecting inflammation-driven enzymatic desialylation, cytoskeletal instability, and metabolic stress within the podocyte.

When comparing different glomerulopathies, distinct glycan signatures emerged. In anti-nephrin– negative podocytopathies, IgM showed a selective reduction in SNA binding, reflecting a loss of terminal sialic acid residues. At the same time, no significant changes were detected with the other lectins tested (AAL, LTL, UEA-I, RCA-I, and ConA), confirming that the alteration was restricted to terminal sialylation. This model supports selective IgM hyposialylation in iNS, inferred from reduced SNA reactivity; orthogonal glycoprofiling will refine site and binding type specificity. In contrast, PLA2R1-positive membranous nephropathy (MN) displayed markedly reduced RCA-I binding, while no significant changes were detected with the other lectins tested (SNA, AAL, LTL, UEA-I, and ConA), indicating that the alteration was restricted to terminal galactosylation. These findings are consistent with those previously reported by Chinello et al. ^27^ for IgG. These opposite glycan patterns underline that antibody-mediated glomerular diseases differ not only by antigen specificity but also by the type of terminal sugar alteration that modulates effector function and target interaction.

Interestingly, in anti-nephrin–positive podocytopathies, our previous study demonstrated that IgG autoantibodies are markedly afucosylated, a modification that enhances FcγRIIIa binding and antibody-dependent cellular cytotoxicity (ADCC) ^5^. Thus, three distinct yet converging forms of immune glycan remodeling can be delineated:

(1) IgM hyposialylation in antibody-negative podocytopathies, altering charge and membrane interactions; (2) IgG hypogalactosylation in PLA2R1-related membranous nephropathy, modifying Fc-mediated effector balance; and (3) IgG afucosylation in anti-nephrin–positive nephrotic syndrome, amplifying FcγR-dependent cytotoxicity.

Together, these observations support an unifying concept in which antibody glycan remodeling defines the effector quality of the humoral response and the modality of podocyte injury.

In antibody-negative nephrotic syndromes, IgM hyposialylation emerges as a structural and charge-dependent factor that directly compromises podocyte integrity, cytoskeletal organization, and energy metabolism, providing a mechanistic basis for antibody-mediated damage in the absence of antigen specificity.

This study has some limitations. Although the in vitro model provided mechanistic insight into how IgM hyposialylation affects podocyte integrity, it cannot fully recapitulate the in vivo environment where complement and circulating mediators also play a role. The use of five pooled IgM preparations reduced interindividual variability but may have masked subtle patient-specific glycan differences. While lectin-based assays demonstrated selective loss of sialic acid, they cannot resolve linkage- or branch-specific changes; LC–MS/MS glycoproteomics and lectin histochemistry on kidney tissue will clarify the precise glycan architecture. The correlation between sialidase abundance, IgM hyposialylation, and proteinuria suggests an enzymatic contribution that should be validated through sialidase inhibition and sialyltransferase activation studies. Finally, longitudinal multi-omic analyses may define how dynamic modulation of IgM sialylation influences podocyte metabolism and clinical outcomes.

Collectively, our results support that IgM hyposialylation, associated with increased serum sialidase levels, is linked to MAPK-driven cytoskeletal remodeling and mitochondrial dysfunction in podocytes. Preservation of a normal phenotype upon enzymatic resialylation provides direct evidence that the terminal sialic acid cap critically regulates the pathogenic potential of circulating IgM. In contrast, IgG afucosylation, as observed in anti-nephrin–positive disease, represents a parallel mechanism of enhanced effector activity, reinforcing the concept that immune glycan remodeling, rather than antigen specificity alone, dictates antibody pathogenicity.

This paradigm also places antibody desialylation within a broader inflammatory context, where the upregulation of NEU1 and NEU3, widely observed in fibrotic and autoimmune diseases, acts as a systemic amplifier of desialylation-induced tissue damage. Targeting the NEU–sialic acid axis may therefore offer a therapeutic opportunity to counteract both the systemic and renal manifestations of immune-mediated damage.

## DISCLOSURE

All authors declare no conflicts of interest.

## DATA SHARING STATEMENT

The mass spectrometry proteomics data have been deposited in the PRIDE repository (ProteomeXchange Consortium) under accession number [**XXX**]. All other data supporting the findings of this study are included in the manuscript and are available from the corresponding author upon reasonable request.

## ACKNOWLEDGMENTS

This work was supported by the Ministero della Salute (Ricerca Corrente), institutional funding from the Department of Experimental Medicine (DIMES), University of Genoa, and the Fondazione Malattie Renali del Bambino ETS.

## Authorship contribution statement

**Sonia Spine**ll**i**: Investigation, Data curation, Writing – original draft. **Andrea Garbarino:** Investigation, Data curation. **Francesca Lugani:** Writing – review & editing, Resources. **Edoardo La Porta:** Writing – review & editing, Resources. **Andrea Petretto:** Writing – review & editing, Investigation, Data curation. **Martina Bartolucci:** Investigation, Data curation. **Chiara Lavarello:** Investigation, Data curation. **Nicole Ginevro:** Investigation, Data curation. **Antonella Trivelli:** Writing – review & editing, Resources. **Giorgio Piaggio:** Writing – review & editing, Resources. **Alberto Magnasco:** Writing – review & editing, Resources. **Maria Ludovica Degl’Innocenti:** Writing – review & editing, Resources. **Sofia Gaudiano:** Investigation, Data curation. Ilaria Musante: Investigation, Data curation. **Paolo Scudieri:** Investigation, Data curation. **Simona Granata:** Writing – review & editing, Data curation. Gianluigi Zaza: Writing – original draft, Resources. **Enrico Verrina:** Writing – review & editing, Resources, Funding acquisition. **Giovanni Candiano:** Writing – original draft, Conceptualization. **Maurizio Bruschi:** Writing – review & editing, Writing – original draft, Supervision, Data curation, Conceptualization, Funding acquisition.

## Notes

### Competing Interest Statement

The authors have declared no competing interest.

