## Supplementary Materials for "IgM hyposialylation drives podocyte injury in pediatric and young adult patients with podocytopathies"

Supplementary Methods

Supplementary Figures 1-3

Supplementary Tables 1-2 (Excel files)

Supplementary Methods

*Subject cohort*

The patients included in this study were the same as those described in our previous paper ^1^, with minor modifications in cohort composition and sample selection. Serum samples from children and young adults affected by iNS with biopsy-proven glomerular disease were obtained from the Unit of Nephrology, Dialysis, and Renal Transplantation at the IRCCS Giannina Gaslini Institute and the Division of Nephrology, Dialysis, and Renal Transplantation at Cosenza (UNICAL). All demographic and clinical characteristics are reported in Table 1. These cohorts included 32 patients with minimal change disease (MCD) and 24 with focal segmental glomerulosclerosis (FSGS). Additionally, 20 patients with primary membranous nephropathy (MN) and 20 with lupus nephritis (LN) were analyzed for comparison. All LN cases were histologically classified as class V, while all MN cases corresponded to stage II. Among the MN patients, 50 % were positive for circulating antibodies against the phospholipase A2 receptor (PLA2R). Samples from 20 healthy donors matched for age and gender were included as additional controls (CTR). According to KDIGO guidelines, nephrotic syndrome was defined by nephrotic-range proteinuria (typically >3.5 g/day in adults or >50 mg/kg/day in children), often accompanied by hypoalbuminemia and edema. For pediatric operational purposes in this study, active disease was defined as an urinary protein-to-creatinine ratio (UPCR) > 2 mg/mg, a threshold commonly used in pediatric nephrology and adopted in previous clinical trials ^2^. Complete remission was defined as a negative or trace protein result on the urine dipstick for three consecutive days, confirmed UPCR of less than 0.2 mg/mg. Relapse was defined by 3+ dipstick for three consecutive days and confirmed by UPCR >2 mg/mg.

For 10 patients with podocytopathies (5 MCD and 5 FSGS), paired serum samples were available at two distinct clinical time points: during active disease (proteinuria > 3.5 g/day) and in complete remission (proteinuria < 0.2 g/day).

Of the 56 patients with FSGS or MCD, 5, 45, and 6 had a clinical phenotype of steroid dependence (SDNS), multidrug dependence (MDNS), and multidrug resistance (MRNS), respectively. More specifically, two consecutive relapses within 15 days of steroid discontinuation defined SDNS; MDNS was defined by the need for two oral immunosuppressive treatments to maintain clinical remission; MRNS was defined by the absence of complete remission after 12 months of treatment with at least two distinct immunosuppressants. Moreover, all FSGS and MCD patients selected were negative for circulating antibodies against nephrin (NPHS1). All subjects or their legal guardians provided written informed consent, and the study was conducted following institutional guidelines and approved by the local ethics committee of the IRCCS Gaslini Institute, Genoa, Italy (Approval numbers 089REG2015) and of the University/Hospital of Foggia, Italy (Approval numbers 191/CE/2023).

*Enzyme-linked immunosorbent assay (ELISA) for IgM N-Glycans*

Nunc MaxiSorp™ ELISA plates (ThermoFisher Cat# 442404) were coated with 100 ng/well of goat anti-human IgM (Merck Cat# I2386) in 50 mM carbonate/bicarbonate buffer (pH 9.6) and incubated overnight (ON) at 4 °C. Uncoated wells served as background controls.

Plates were washed three times with PBS, blocked ON at 4 °C with SuperBlock (ThermoFisher Cat# 37518), rewashed, and incubated ON at 4 °C with 100 μL of 1:100 diluted serum, standard, or positive/negative controls in SuperBlock + 0.05 % Tween (SuperT).

After five washes, 100 μL of a single biotinylated lectin was added to each well: Sambucus nigra agglutinin (SNA; Vector Laboratories Cat# B-1305-2), preferentially binding terminal α2,6-linked sialic acid; Aleuria aurantia lectin (AAL; Cat# B-1395-1), recognizing fucose residues with preference for α1,6-linked core fucose; Ulex europaeus agglutinin I (UEA-I; Cat# B-1065-2), binding mainly α1,2-linked fucose; Lotus tetragonolobus lectin (LTL; Cat# B-1325-2), showing affinity for α1,3-linked fucose (Lewis-type structures); Ricinus communis agglutinin I (RCA-I; Cat# B-1085-1), preferentially recognizing terminal β-galactose residues; and Concanavalin A (ConA; Cat# B-1005-5), binding mannose residue typical of high-mannose or hybrid N-glycans. Lectins, diluted 1:1000 in SuperT, were added to each well, and incubated ON at 4 °C. Plates were washed, incubated with 100 μL of streptavidin-HRP (ThermoFisher Cat# 434301, 1:10,000 in SuperT) for 30 min at room temperature (RT), rewashed, and developed with 100 μL of TMB substrate (Bio-Rad Cat# 1721066).

The reaction was stopped with 100 μL of 100 mM H₂SO₄, and OD₄₅₀ was measured using a Bio-Rad iMark reader (Cat# 1681130).

N-glycan residue levels were determined using a standard curve obtained from serial dilutions of the highest-titer healthy donor serum. For each sample, lectin reactivity was normalized to the corresponding total IgM levels, measured in parallel by anti-IgM ELISA (ThermoFisher; Cat# BMS2098), to account for interindividual differences in IgM concentration. The detection limit of each N-glycan residue corresponded to the lowest value distinguishable from blank.

Lectin specificity was verified by pre-incubating 100 μg of purified serum IgM diluted in PBS and treated with N-glycosidase F (PNGase F; New England Biolabs Cat# P0704, 1 U/μg protein) ON at 37 °C to remove N-glycans ^3^. Samples were then concentrated and washed using Amicon Ultra filters (Merck Cat# UFC9100) to remove N-glycans and enzymes. Treated samples were reanalyzed with lectin-based ELISA under identical conditions.

N-glycan residue levels were reported as median ± interquartile range and visualized as box plots.

*Enzyme-linked immunosorbent assay (ELISA) for sialytransferase- and sialyhydrolase-related enzymes*

Nunc MaxiSorp™ ELISA plates (ThermoFisher Cat# 442404) were coated with 60 µg/well of serum sample diluted in 50 mM carbonate/bicarbonate buffer (pH 9.6) and incubated ON at 4 ^◦^C. Uncoated wells served as background controls. Plates were washed three times with PBS, blocked ON at 4 ^◦^C with SuperT (ThermoFisher Cat# 37518), rewashed, and incubated ON at 4°C with 100 μL of rabbit anti-Human ST6 beta-galactoside alpha-2,6-sialyltransferase 1 (ST6GAL1; ThermoFisher; Cat# 14355-1-AP), or rabbit anti-Human neuraminidase 1 (NEU1; Merck; Cat# HPA015634) or rabbit anti-Human neuraminidase 3 (NEU3; Merck; Cat# HPA070381) diluted 1:1000 SuperT. After five washes, 100 μL of goat anti-Rabbit HRP-conjugated (R&D system; Cat# HAF008) diluted 1:2000 in SuperT was added and incubated for 2 h at RT. Plates were rewashed and developed with 100 μL of TMB substrate (Bio-Rad Cat# 1721066). The reaction was stopped with 100 μL of 100 mM H_2_SO_4_. OD_450_ was measured using a Bio-Rad iMark reader (Cat# 1681130). Enzyme levels were determined using a standard curve from serial dilutions of the highest-titer healthy donor serum. The enzyme detection limit corresponded to the lowest value distinguishable from the blank. Enzyme levels were reported as median and interquartile range and visualized as box plots.

*IgM purification*

Serum IgM was purified using a 1 mL HiTrap™ IgM Purification HP column (Cytiva, Cat# GE90100422) according to the manufacturer’s instructions. Briefly, five independent iNS IgM pools, each obtained by combining sera from four distinct patients with low sialic acid levels and comparable SNA/anti-IgM ratios, and five control (CTR) pools, each prepared from four age- and sex-matched healthy donors, were processed in parallel.

For each pool, 1 mL of serum was diluted with binding buffer (20 mM sodium phosphate, 0.8 M (NH₄)₂SO₄, pH 7.5) and loaded onto a column pre-equilibrated with the same buffer.

The column was washed with 10 column volumes (CV) of binding buffer to remove unbound material, and IgM was eluted with 5 CV of elution buffer (20 mM sodium phosphate, pH 7.5). After elution, the column was regenerated with 10 CV of binding buffer. Eluted IgM fractions were concentrated and washed in PBS using Amicon Ultra-15 centrifugal filters with a 100 kDa molecular weight cut-off (Merck, Cat# UFC910008) to remove salts and low-molecular-weight contaminants.

The final preparations were filtered through 0.45 μm membranes under sterile conditions before use in podocyte experiments.

SDS-PAGE verified the purity and integrity of all IgM preparations under reducing conditions and visualized by Blue Silver staining.

*Desialylation and resialylation of CTR human IgM*

Human IgM purified from pooled CTR serum was desialylated by incubation with neuraminidase (Merck, Cat# 10269611001) at 37 °C for ON in 50 mM sodium acetate buffer (pH 6.0) under gentle agitation. One mU of enzyme per 100 µg of IgM was used in a final reaction volume of 100 µL. After incubation, IgM was concentrated and washed in PBS using Amicon Ultra-15 centrifugal filters with a 100 kDa molecular weight cut-off (Merck, Cat# UFC910008) to remove salts, enzymes, and low-molecular-weight contaminants. SNA lectin ELISA assessed desialylation efficiency.

For resialylation, desialylated IgM was incubated with β-galactoside α2,6-sialyltransferase 1 (ST6GAL1; Creative Biomart, Cat# ST6GAL1-28H) in the presence of cytidine 5′ monophospho-N-acetyl neuraminic acid (CMP-Neu5Ac) (Merck, Cat# ADVH9A98841C) as donor substrate.

The reaction was performed in 50 mM Mops-HCl buffer (pH 7.5) containing 10 mM MnCl₂, using 1 mU of enzyme per 100 µg of IgM and 20 mM CMP-Neu5Ac, for 48 h at 37 °C under gentle agitation in a dark box ^4^. Following the reaction, salts, enzymes, and residual donor substrate were removed by ultrafiltration through the same Amicon filters. SNA lectin ELISA confirmed the successful restoration of sialic acid residues.

*Cell culture and IgM treatment*

Podocyte cell culture, as previously described and characterized ^5, 6^, was grown in Dulbecco’s modified Eagle’s medium (DMEM) high glucose supplemented with 10% FBS, 2 mM L-glutamine, penicillin (100 U/mL), and streptomycin (100 μg/mL). Before use, all sera were heat-inactivated at 56 °C for 45 min to ensure complement inactivation. Cells were seeded at subconfluent density and then exposed ON to IgM from iNS or CTR, either unmodified or enzymatically treated to yield desialylated (Desialylated) or resialylated (Resialylated) forms. All experimental procedures were conducted under aseptic conditions to exclude contamination, and the occurrence of nonspecific pyrogenic effects is considered highly unlikely.

*Confocal microscopy*

Immunofluorescence (IF) studies were performed on podocytes cultured on glass slides and treated with IgM iNS or IgM CTR, either without modification, desialylated, or resialylated. Briefly, cells were fixed with 4% paraformaldehyde in PBS for 15 min at RT. After 3 washes in PBS, cells were permeabilized with 0.5% Triton X-100 (BioRad; Cat# 1610407) in PBS for 10 min, blocked with 3% BSA in PBS for 1 h at RT, and then incubated 1h with phalloidin–Alexa Fluor™ 488 (Invitrogen Cat# A12379) or ON at 4 °C with sheep anti-human nephrin antibody (R&D Systems Cat# AF4269, 1 μg/mL), followed by donkey anti-sheep IgG–Alexa Fluor™ 568 (Invitrogen Cat# A-11015) for 1,5 h at RT. All Alexa Fluor™ conjugates were used at 1:500 in a humid, dark chamber. After 3 washes in PBS, the chambers were removed, and the slides were mounted with Fluoroshield with DAPI to stain cell nuclei. Imaging was performed using a laser scanning confocal microscope TCS SP8 (Leica Microsystems). Images were analyzed using the EBImage package version 4.51.0 ^7^.

*Mass Spectrometry*

Podocyte cells were lysed, reduced, and alkylated in 50 μL LYSE buffer (Preomics) at 95°C for 10 min and sonicated with an Ultrasonic Processor UP200St (Hielscher), 3 cycles of 30 sec. Proteins were isolated and digested by the PAC method, automated on a KingFisher™ Apex robot (Thermo Fisher Scientific) in 96-well format according to Bekker-Jensen et al ^8^. Briefly, the tip plate was stored in plate #1. Lysate samples were stored in plate #2, in a final concentration of 70% acetonitrile and with magnetic beads in a protein/bead ratio of 1:4 (1:1 SpeedBead Magnetic Carboxylate, 45152105050250 and 65152105050250). Washing solutions were in plates #3–5 (acetonitrile), plate #6 (70% Ethanol), and plate #7 (isopropanol). Plate #8 contained 100 μL digestion solution of 25 mM Tris HCl pH 8, LysC (Wako) in an enzyme/protein ratio of 1:100 (w/w) and trypsin (Promega) in an enzyme: protein ratio of 1:50. The protein aggregation was carried out in two steps of 1 min mixing at medium mixing speed, followed by a 10 min pause each. The sequential washes were performed in 2.5 min at a slow speed, without releasing the beads from the magnet. The digestion was set to 4 h at 37 °C with slow speed.

The resulting peptides were analyzed by a nano-UHPLC-MS/MS system using an Ultimate 3000 RSLC coupled to an Orbitrap Q Exactive Plus mass spectrometer (Thermo Scientific Instrument). Elution was performed with an EASY spray column (75 μm × 50 cm, 2 μm particle size, Thermo Scientific) at a flow rate of 250 nL/min using a linear gradient of 2-45% solution B (80% acetonitrile, 5% dimethylsulfoxide, 0.1% formic acid in H_2_O) in 50 min. Orbitrap detection was used for MS1 measurements at a resolving power of 70 K in a range between 375 and 1500 m/z with a 3x10^6 automatic gain control (AGC) target and 50 ms maximum injection time (IT). Precursors were selected for data-independent fragmentation with an isolation window width of 34 m/z and 19 loop count. Higher collisional dissociation (HCD) energy was set to 27%, and MS2 scans were acquired at a resolution of 35 k, 3x10^6 AGC target, and 50 ms maximum IT.

All DIA raw files were processed with Spectronaut software version 18 ^9^ using a library-free approach (directDIA) under default settings. The library was generated against the Uniprot Human database (release UP000005640_9606 June 2024, 104602 Entries). Carbamidomethylation was selected as a fixed modification, while methionine oxidation and N-terminal acetylation were selected as variable modifications. The false discovery rate (FDR) for peptide-spectrum matches (PSMs) and peptide/protein groups was set to 0.01. For quantification, Precursor Filtering was set to Identified (Qvalue), and MS2 was chosen as the quantity MS level.

*Kinome signaling*

To evaluate the phosphorylation signaling in cultured podocytes treated with IgM iNS, IgM CTR, or desialylated/resialylated CTR IgM, we used the Human Phosphorylation Pathway Profiling Array C55 kit (Raybiotech, Cat# AAH-PPP-1-2) following the manufacturer’s instructions. Briefly, the array was incubated with blocking buffer at RT for 30 min, then incubated with 100 μg of cell lysate ON at 4 °C. The array was washed twice with wash buffer, then incubated with the Detection Antibody Cocktail at RT for 2 h. After washing, the array was incubated with horseradish peroxidase-conjugated secondary antibody at RT for 2 h. Images were acquired with the ChemiDoc Imaging System (Bio-Rad Cat# 12003153) and analyzed using the KSEA App ^10^. Each experiment was performed in triplicate.

*Lipid peroxidation assay*

Lipid peroxidation was assessed by quantifying malondialdehyde (MDA) levels using the Malondialdehyde (MDA) Colorimetric Assay Kit (TBA method, Thermo Fisher Scientific, Cat. No. EEA015) according to the manufacturer’s instructions. The assay is based on the reaction between MDA and thiobarbituric acid (TBA) to form a red adduct with maximal absorbance at 532 nm, reflecting the extent of lipid peroxidation. Podocytes treated with IgM iNS, IgM CTR, or desialylated/resialylated CTR IgM were lysed in ice-cold PBS, and the homogenates were centrifuged at 10,000 × g for 10 min at 4 °C. The resulting supernatants were mixed with the acid reagent and chromogenic agent provided in the kit, and incubated at 95 °C for 40 min to allow color development. After cooling and centrifugation (10,000 × g for 10 min), 250 μL of each supernatant was transferred to a 96-well microplate, and absorbance was measured at 532 nm using a Spark multimode microplate reader (TECAN, Cat# 399950). A standard curve was generated using MDA standards. MDA concentration in each sample was expressed as nmol MDA per mg of total protein, and protein content was determined using the Quick Start Bradford Assay (Bio-Rad; Cat. No. 5000205). Each condition was analyzed in five independent biological replicates.

*Measurement of adenosine triphosphate (ATP) synthesis*

Intracellular ATP levels were quantified using the ATP Determination Kit (ThermoFisher; Cat# A22066), based on the luciferin–luciferase bioluminescence method, according to the manufacturer’s instructions. This assay relies on the ATP-dependent oxidation of D-luciferin by recombinant firefly luciferase, producing a light signal with a maximum emission at ~560 nm, directly proportional to the ATP concentration. Podocytes treated with IgM iNS, IgM CTR, or desialylated/resialylated CTR IgM were lysed in ice-cold PBS and centrifuged at 10,000 × g for 10 min at 4 °C to remove debris. The supernatants were mixed with the luciferase reaction mixture containing 0.5 mM D-luciferin, firefly luciferase (1.25 µg/mL), 5 mM MgSO₄, 100 µM EDTA, 1 mM DTT, and 25 mM Tricine buffer at pH 7.8. Luminescence was measured immediately using a Spark multimode microplate reader (TECAN, Cat# 399950).

A standard curve was generated using ATP standards, and the levels were expressed as nmol ATP per mg of total protein, with protein concentration determined by the Quick Start Bradford Assay (Bio-Rad; Cat# 5000205). Each condition was analyzed in fine independent biological replicates.

*Bioinformatic and statistical analyses*

After log2 conversion, a normal distribution was used to impute missing values, and the entire dataset was normalized using the quantile method. The normalized dataset was analyzed using unsupervised hierarchical clustering, including principal component analysis and k-means clustering, to identify outliers and dissimilarities among samples. ANOVA test for unpaired samples was used to identify statistically significant proteins between podocyte cells treated with iNS IgM-derived (iNS), CTR IgM-derived, without modification (CTR) or desialylated (Desialylated). Then, to identify the proteins that maximized discrimination between iNS and CTR, or Desialylated and CTR, or Desialylated and iNS, we used t-test analysis. For both analyses, proteins were considered statistically significantly expressed for power values of 80% and adjusted p-values ≤ 0.05 after correction for multiple interactions (Benjamini-Hochberg method). Volcano plots were used to visualize the results of the t-test analysis, with the threshold line determined by the function y = |c/(x − x0)|. Gene ontology analysis was performed to identify the enriched pathways through the active subnetworks using the PathFindR package ^11^. Kinase enrichment analysis (KEA) was also performed using the statistically significant proteins as substrates to predict kinases potentially involved in regulating the phosphorylation signal in each comparison ^10^. The Coral App was used to visualize the kinome tree of proteome profile ^12^.

For the lectin ELISA, lipid peroxidation, and ATP synthesis differences across all clinical groups were determined using the Kruskal-Wallis test with Dunn's correction. Pearson’s correlation coefficients were calculated to assess the relationships between UPCR, SNA reactivity, and serum NEU1 and NEU3 levels. Paired samples collected at <0.2 and >3.5 gr/day proteinuria were compared using the Wilcoxon test. For the kinome assay, differences across all conditions were determined using a t-test with Benjamini-Hochberg correction. All statistical tests were performed using OriginLab Pro version 2022b and R version 4.3.3 software.

Supplementary Figure S1

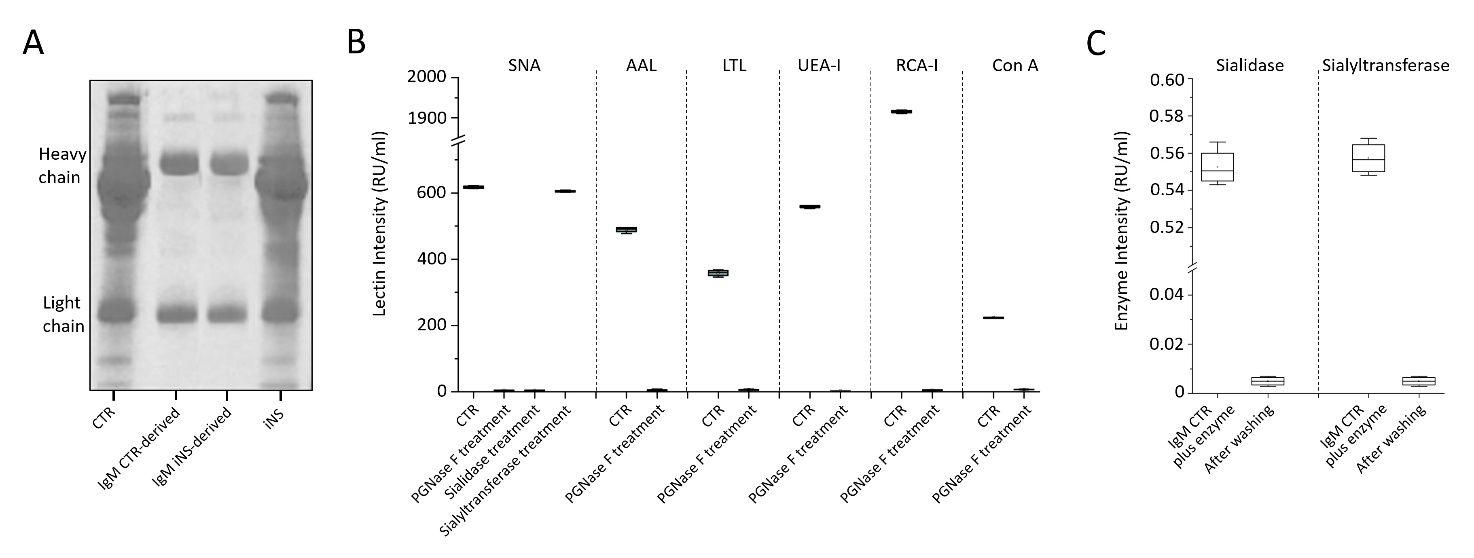

Figure S1. Validation of IgM purity, enzymatic desialylation/resialylation efficiency, absence of residual enzyme activity, and lectin specificity.

(A) Validation of IgM purity. SDS-PAGE of purified IgM under non-reducing conditions stained with Blue Silver, confirming high purity of the preparations. (B) Validation of lectin specificity and enzymatic treatments. Purified serum IgM was pretreated with PNGase F to remove N-linked glycans prior to lectin ELISA, which completely abolished lectin binding and confirmed that signals reflect specific N-glycan–dependent recognition. SNA lectin ELISA further demonstrated a complete loss of terminal sialic acid binding after neuraminidase treatment (Desialylated IgM), comparable to PNGase F-treated IgM, and reestablishment of SNA binding to levels comparable to untreated IgM after resialylation (Resialylated IgM). (C) Validation of enzyme removal from purified IgM. ELISA specific for sialidase or sialyltransferase performed on desialylated and resialylated IgM confirmed the absence of enzyme carryover, ensuring that podocyte responses were not affected by residual enzymatic activity.

Supplementary Figure S2

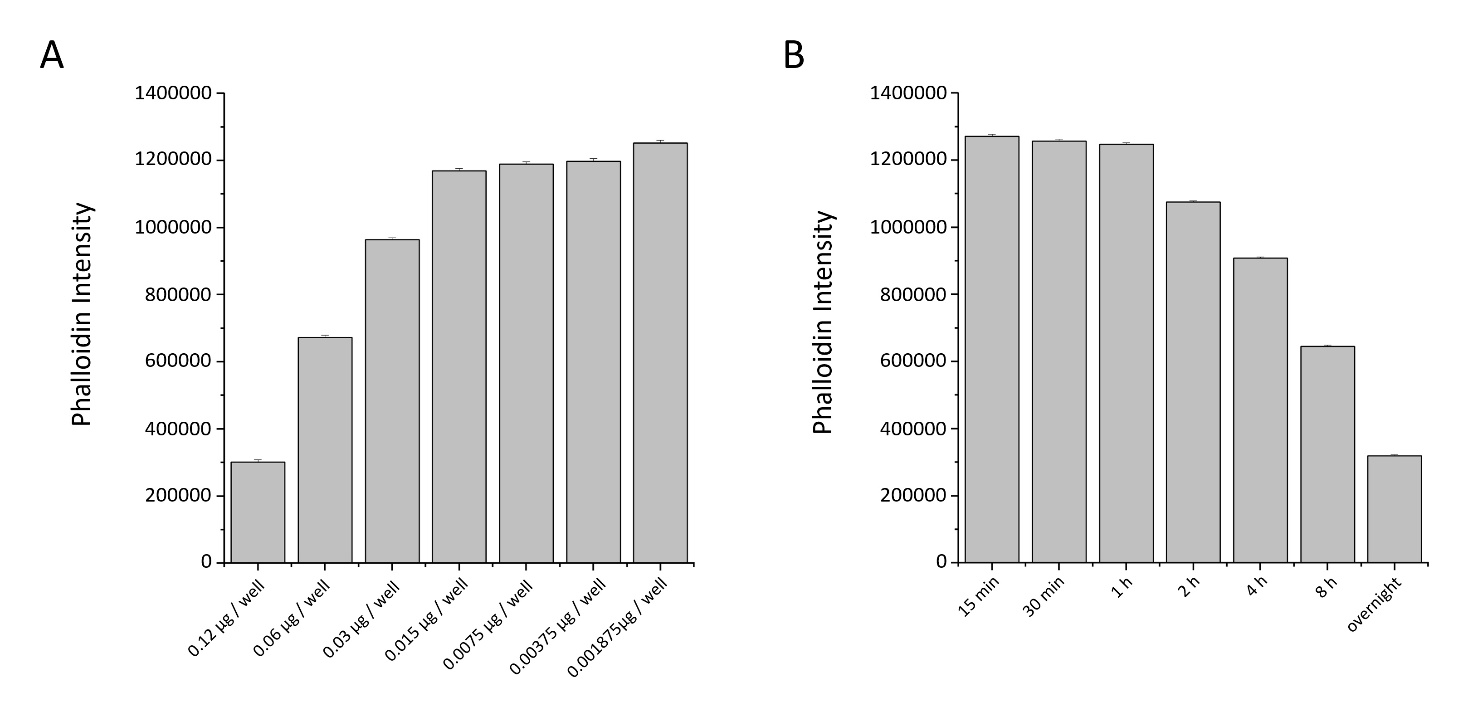

Figure S2. Optimization of purified IgM treatment conditions on human podocytes.

Podocytes were incubated with purified IgM at concentrations ranging from 0.12 to 0.0019 µg per well (corresponding to serial dilutions equivalent to whole serum from 1:50 to 1:4000). Incubation times varied from 15 min to overnight (ON). Quantitative analysis of actin organization, normalized to the number of cells, was performed using automated image processing with the EBImage package in R. Histograms show the total phalloidin fluorescence intensity ± standard deviation and reveal a direct correlation between IgM concentration (A) and incubation time (B) with the degree of cytoskeletal rearrangement. The 0.12 µg IgM/well under overnight incubation was selected as the standard condition, producing the maximal and most reproducible effect.

Supplementary Figure S3

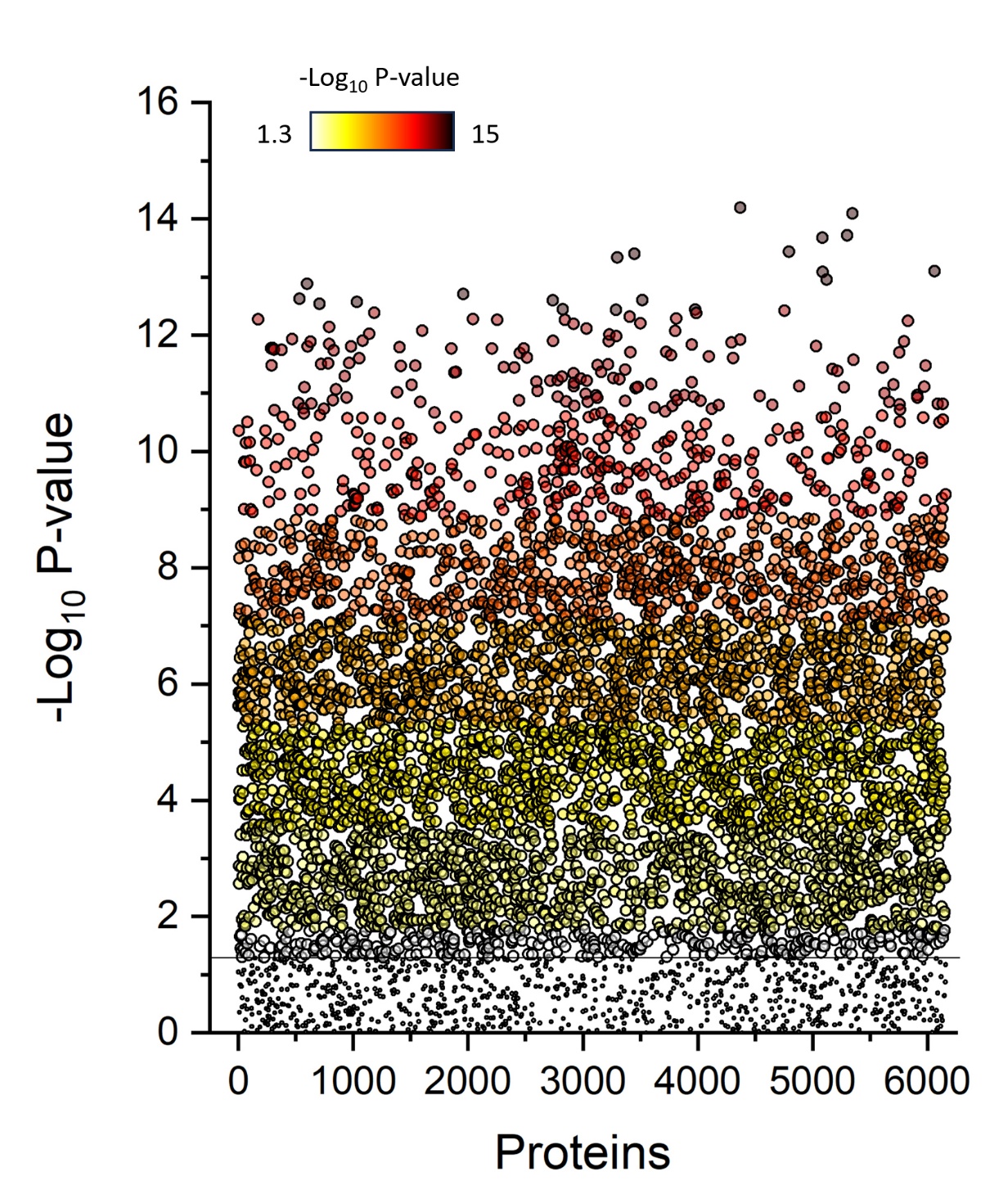

Figure S3. Two-dimensional plot of ANOVA-significantly modulated proteins in podocytes.

Two-dimensional plot of 5,250 ANOVA-significantly modulated proteins. Each circle represents a protein, color-coded according to statistical significance (white = –log₁₀ 1.3; yellow → orange → red → black = –log₁₀ 15). The black line indicates the –log₁₀ 1.3 significance threshold.

Supplementary Figure 4

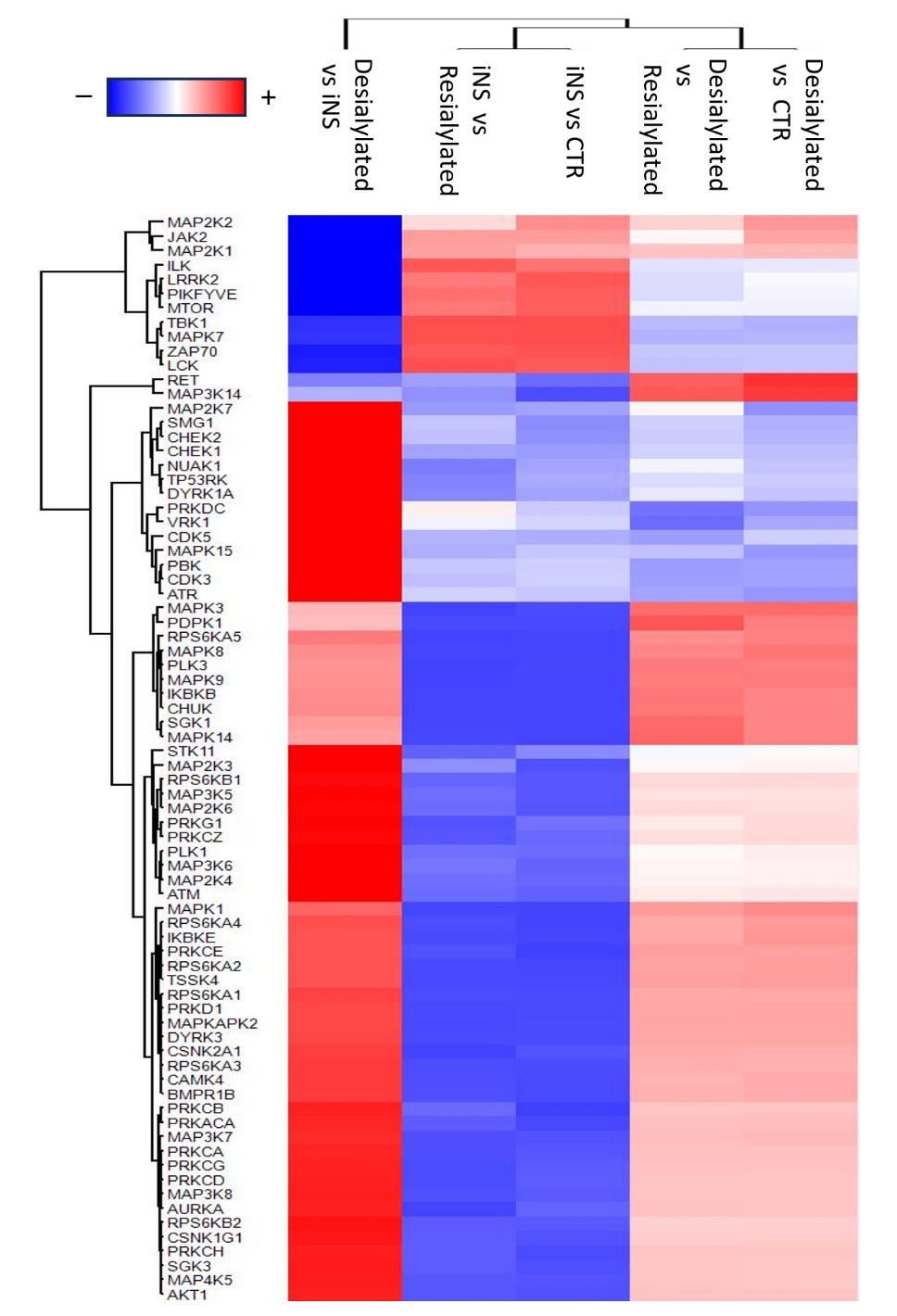

Figure S4 Kinome enrichment analysis.

Heatmap of kinase activity inferred by Kinase-Substrate Enrichment Analysis (KSEA) of phospho-array data. Each row represents a kinase, and columns correspond to podocytes treated with CTR, iNS, Desialylated, or Resialylated CTR IgM. Color intensity indicates relative kinase enrichment scores. Desialylated IgM treatment markedly enhanced MAPK (ERK1/2, p38, JNK) and SRC pathway activation while suppressing AKT/mTOR-associated kinases. Resialylation maintains the kinase activity profile to levels indistinguishable from control.

Table S1. List of significantly modulated proteins identified by quantitative proteomics.

Comprehensive dataset of differentially expressed proteins identified across all comparisons. For each entry, UniProt accession, gene name, log₂ fold change, and adjusted p-value (Benjamini–Hochberg correction) are reported.

Table S2. Gene Ontology (GO) enrichment analysis of significantly modulated proteins.

List of 38 significantly enriched GO Biological Process terms obtained from PathFindR analysis of the proteomic dataset. Each term includes GO ID, term description, fold enrichment score, Log10 p-value (FDR correction), and number of associated genes. GO terms could be grouped into five main clusters: MAPK/stress signaling, cytoskeletal and adhesion processes, mitochondrial metabolism, oxidative stress responses, and inflammatory signaling.
